# YBX2 Dysregulation in Maturation Arrest NOA YBX2 Dysregulation is a Potential Cause for Late Maturation Arrest in Men with Non-Obstructive Azoospermia

**DOI:** 10.1101/364968

**Authors:** Ryan Flannigan, Anna Mielnik, Brian D. Robinson, Francesca Khani, Alex Bolyakov, Peter N. Schlegel, Darius A Paduch

## Abstract

**Introduction:** YBX2 protein binds to Y-box promotors and to mRNAs in the cytoplasm of pachytene spermatocytes and round spermatids in rodents. Knock-out of YBX2 leads to maturation arrest in animal models. YBX2 binds PRM1 and 2 mRNA, which is transcribed early and sequestrated for translation during late spermatogenesis.

**Objective:** This study aimed to determine if the loss of YBX2 is associated with MA arrest due to the loss of sequestration of protamines in the human testis. As a second aim, we examined the expression of YBX2, and its transcription factors in maturation arrest (MA)(early and late) and normal controls in men.

**Methods:** RNAseq was performed using RNA extracted from human testis samples from 44 men with non-obstructive azoospermia and ten from healthy controls. Differential expression was performed using JMPgenomics, FDR<0.001. FANTOM5 was used to predict enhancers and inhibitors of YBX2 expression. Immunofluorescence (IF) was used to stain testis tissue sections with antibodies against YBX2, SYCP3, and PRM2 in normal and MA samples. Flow cytometry was utilized to characterize YBX2 positive cells.

**Results:** Expression of *YBX2* mRNA was significantly downregulated in early and late MA compared to controls. Surprisingly, PRM1&2 mRNAs were also depleted in men with MA. Multifactorial regression analysis demonstrated a decrease in YBX2 expression in MA is due to decrease in COMP levels (p<0.0001). IF localized YBX2 protein in spermatocytes and round spermatids among fertile men, with rare YBX2 positive spermatocytes stained in LMA. PRM1&2 proteins were absent or abnormally sequestrated within spermatocytes.

**Conclusions:** Decrease in YBX2 protein expression in men with LMA leads to loss of translational suppression and lack of PRM1 and PRM2 necessary to complete spermatogenesis.

## Introduction

The most severe form of male infertility is azoospermia and occurs in 10-20% of men presenting with infertility^1^. In non-obstructive azoospermia (NOA), the abnormal progression of spermatogenesis results in a lack of mature sperm. Histological abnormalities during spermatogenesis result in early (eMA) or late maturation arrest (lMA) at spermatocyte stage or spermatid stage respectively, or even earlier with absent or limited spermatogonial stem cells (SSCs) termed Sertoli cell-only syndrome (SCO). Errors in spermatogenesis may occur during SSCs renewal and differentiation (mitosis), meiosis 1 or 2, or during post-meiotic spermiogenesis. Our work has focused on understanding how small RNAs regulate expression of mRNAs during spermatogenesis. This manuscript answers if regulation of YBX2 leads to downstream reproductive effects, with separate paper focused on miRNAs and YBX family interactions.

Before completion of spermiogenesis, histones are replaced by protamines for chromatin condensation to take place. Transcription of messenger RNAs (mRNA) and its translation to proteins is temporally uncoupled during most stages of meiosis ^2–5^. RNAs are sequestrated from translational machinery via RNA binding proteins (RBPs). Y-Box family RNA binding proteins are necessary for completion of spermatogenesis and thus fertility. In mice, YBX2 (MSY2) is highly abundant in spermatids, accounting for 0.7% of all protein present among these cells^6^. MSY2 knock outs results in infertility with maturation arrest phenotype in transgenic mice. MSY2 has dual functions in rodents; it can act as a suppressor of mRNA translation by binding to mRNAs in meiotic cells and round spermatids^7^. MSY2 can also function as a nuclear transcription factor enhancing transcription of genes essential in the completion of spermatogenesis such as protamine 2^8,9^. Timely translation of PRM1/2 is vital for chromatin modeling in elongating spermatids and exchange of histones to protamines ^2–5^. MSY2 also functions as a nuclear shuttle protein removing transcripts from the nucleus among spermatocytes^6,10,11^.

Although YBX2 knockout is sterile due to maturation arrest in mice, initial observations by Yang has not subsequently been extensively studied in the human testis. In humans, YBX2 has a testis-specific expression; its gene localizes to chromosome 17p11.2-13.1^12^. Among infertile men with NOA, YBX2 mRNA expression, tested by RT-PCR, was reduced in testes samples compared to fertile controls^13^. Hammoud *et al*. (2009) discovered a higher rate of polymorphisms of YBX2 in men with NOA, identifying 15 polymorphic sites with 7 of SNPs reaching statistical significance^14^. Two of these polymorphisms led to amino acid substitutions in the cold shock domain indicating that YBX2 polymorphisms are associated with NOA. A recent study evaluating infertile men defined as those unable to conceive for at least 12 months did not have increased rates of polymorphisms of YBX2 compared to fertile controls.

YBX2 plays an essential role in late stages of spermatogenesis by regulating the timing of translation in spermatocytes and round spermatids. Mutations in the cold shock domain of YBX2 are associated with NOA. However, the specific function of YBX2, including its dynamic expression and translation, as well as the particular RNAs bound and regulated by YBX2 are mostly unknown among infertile men, specifically among those with maturation arrest NOA. We hypothesize that dysregulation of YBX2 expression during spermatogenesis leads to the loss of essential mRNAs (i.e., PRM1 and 2) and proteins to complete spermatogenesis. In this study, we aimed to evaluate the expression of YBX2, its predicted target mRNAs, and YBX2 promoter regulators using next-generation RNA sequencing (NGS) in a large number of men with NOA and different histologies. Immunofluorescence and flow cytometry were used for localization and enumeration of YBX2, PRM2, and SYCP3 proteins in both healthy controls and among men with MA to gain insight with respect to how aberrant expression of YBX2 may lead to maturation arrest.

## Methodology

This study was performed under IRB approval from Weill Cornell Medicine (WCM), New York, NY.

### Subjects

Testis tissue was extracted from 44 men with NOA and ten fertile controls. Among the 44 patients with NOA, 11 had SCO, 11 had eMA, 5 had lMA, 10 had HS, and 7 had KS. Testis biopsies were harvested from men with NOA at the time of microdissection testicular sperm extraction (mTESE). A small portion of the tissue was placed in Bouins fixative, stained with hematoxylin and eosin (H & E) and examined by light microscopy. Histology was performed by a genitourinary pathologist at WCM and classified as SCOS, eMA, lMA, or HS. As SCO testes lack meiotic germ cells, there were used for negative controls. For controls, testis tissue was harvested from cadaveric organ donors with normal spermatogenesis. Normal testis histology was defined by seminiferous tubules comprised of thin basement membranes, normal germinal epithelium and orderly progression of maturing germ cells: spermatogonia, spermatocytes, spermatids and mature spermatozoa^15^. The remainder of the tissue was snap frozen in Eppendorf tubes and used for RNA isolation and sequencing.

### RNA Isolation and Quality Control

All procured tissues were snap frozen in liquid nitrogen. Total RNA was isolated from all testis tissue using miRCURY RNA Isolation Kit (Cat No.300111, Exiqon Inc. Vadbeak, Denmark). First, the frozen tissue was completely homogenized in a lysine solution using TissueRuptor. The lysate was incubated with proteinase K for protein removal and loaded on a spin column. Genomic DNA was removed directly on the column with DNase I (Qiagen) at final concentration of 0.25 Kunitz unit/microliter during the 20-minute incubation. Eluted total RNA was checked for purity and integrity on Agilent Bioanalyzer 2100 (Agilent Technologies, CA, USA). Only RNA’s with RNA Integrity number (RIN) greater or equal to 7 that not show any RNA degradation were used. RNA sample quality was confirmed by Nanodrop spectrophotometer to determine the purity based on ratio A260/280 indicative of protein contamination and A260/A230 indicative of chemical contamination of salts, phenol, and carbohydrates. RNA concentration was measured with a fluorescence based quantitation assay using Q-bit fluorometer (Life Technology, NY, USA).

### Illumina Library Preparation

TruSeq-barcoded RNAseq libraries were generated with the NEBNext Ultra RNA Library Prep Kit (New England Biolabs). Each library was quantified with a Qubit 2.0 (dsDNA HS kit; Thermo Fisher), and the library sizes distribution was determined with a Fragment Analyzer (Advanced Analytical) before pooling.

### Illumina Sequencing

Libraries were sequenced using an Illumina NextSeq500. At least 20M single-end reads with a minimum length of 75 base pairs were generated per library. *Preprocessing*: reads were trimmed for low quality and adaptor sequences with cutadapt. Parameters included: -m 10 -q 20 -a AGATCGGAAGAGCACACGTCTGAACTCCAG –match-read-wildcards *Genome mapping:* reads were collapsed and mapped to the reference genome (UCSC hg19) using Tophat and Star aligners. Cufflinks were used for feature extraction, and gene counts tables preparation.

### Statistical Analysis

Statistical analysis was performed using JMP Pro version 12. FDR<0.001 was used to identify differentially expressed genes.

### Identification of YBX2 transcription factors using RNAse and FANTOM5 database

FANTOM5 human promoterone view Hg19 was used to identify predicted promotors of YBX2^16^. This pipeline has been previously described^16^. In brief, FANTOM5 draws on a compiled dataset created through tissue-specific cap analysis of gene expression (CAGE) to identify and map transcription starting sites and transcription factors such as activators or repressors. The identified promotors for YBX2 were then evaluated from our dataset using a multifactorial regression analysis to determine if the aberrant expression of promotor regulators leads to dysregulation of YBX2.

### Immunofluorescence

Testis tissue was formalin fixed, and paraffin embedded, and sectioned to the 5 um thickness and mounted onto glass slides. Slides were de-waxed using three washes of 100% xylene, followed by serial rehydration in ethanol baths. Epitope retrieval was performed using Dako target (pH9) retrieval solution, for 30 minutes in a 96°C water bath. Sections were permeabilized using phosphate buffered saline with Triton (PBST) 0.2% for 30 minutes, following by blocking with 1x phosphate buffered saline (PBS), 5% Normal Goat Serum, 3% bovine serum albumin (BSA), 0.2% Triton for 90 minutes. Primary antibodies were used to stain YBX2 (MSY2 mouse antibody Novus Bio H0005087; 1:100), SYCP3 (rabbit Abcam ab15093; 1:100), and PRM2 (rabbit NBP2-31637; 1:100). Slides were incubated with the primary antibodies at room temperature for 2 hours. Following washing of the primary antibody, secondary antibodies (AF488, and AF555) were applied at a concentration of 1:1000 for 60 minutes at room temperature in the dark. Prolong Gold antifade with DAPI was added to the slide along with a glass coverslip. Negative controls (no primary antibody) were acquired and processed utilizing the same parameters. The stains were performed at least three times for each antibody.

### Fluorescent Microscopy

Images were acquired using a Nikon Eclipse 50i system, illuminated with the Nikon Intensilight Hg Pre-centered Fiber Illuminator C-HGFIE. The Nikon DSQi2 monochrome immunofluorescent camera was used to acquire each channel. Z-stacks were utilized for deconvolution (z-step=0.06 um and 96.5 nm per pixel). NIS- Elements in Basic Research, version 3.10, SP3, Nikon and FIJI for Mac was used to process the flat images and deconvolute the z-stacks utilizing DeconvolutionLab2 plugin respectively. All images for the respective antibodies were acquired with the same gain and exposure times. Cell immunofluorescence intensity and size were calculated using FIJI software for Mac.

### Flow Cytometry

Testis tissues were mechanically disintegrated using the MediMachine (BD Biosciences, San Jose CA, USA). The cells were filtered with 50um and 30um mesh to remove cellular clumps and extracellular matrix. Cells were then fixed using CytoFix-Cytoperm (BD Biosciences, San Jose CA, USA), for 20 minutes. Specimens were then centrifuged at 600 relative centrifugal force (RCF) for 5 minutes and washed in a 1x Perm-wash solution (BD Biosciences, San Jose CA, USA), through 2 additional 5-minute centrifugation steps, and a 10 minute incubation period. Primary YBX2 (MSY2 mouse antibody Novus Bio H0005087; 1:100) antibody was added and incubated for 1 hour on ice. The cells were washed with PBS-BSA solution with two centrifugation steps for 5 minutes at 500 RCF. Secondary anti-mouse AlexaFluor 555 (1:500 dilution) antibody was added for a 30-minute incubation period in the dark, on ice.

Stained samples and F-1 controls were then run on the BD Acurri C6 (BD Biosciences, San Jose CA, USA) flow cytometer. The laser filter to detect AF555 included 585/40 in FL2. Samples were processed until 100,000 events or until the entire 300ul sample was processed. Water and unstained samples were used as controls. FlowJo version 10.1r1 (Ashland Oregon, USA) was used to analyze FCS files. 7-AAD with forward and side scatter were used to identify 1N, 2N, and 4N populations within the testis. FSCH/FSCA gates were applied to eliminate duplets. The same gates were used for all experiments. Compensation was performed by using single antibody samples. YBX2 positive cell populations demonstrating positive fluorescence in FL2 were compared to unstained cells and SCO samples which lack meiotic cells. All experiments were repeated at least three times.

## Results

Results of RNA sequencing analysis revealed that amount of *YBX2 mRNA* was very low in SCO as expected since there are no meiotic germ cells in testes with SCO. YBX2 mRNA was abundantly present in early and late maturation arrest, and normal spermatogenesis samples. Thus, indicating that *YBX2* is transcribed in meiotic germ cells, Figure 1.

**Figure 1.**
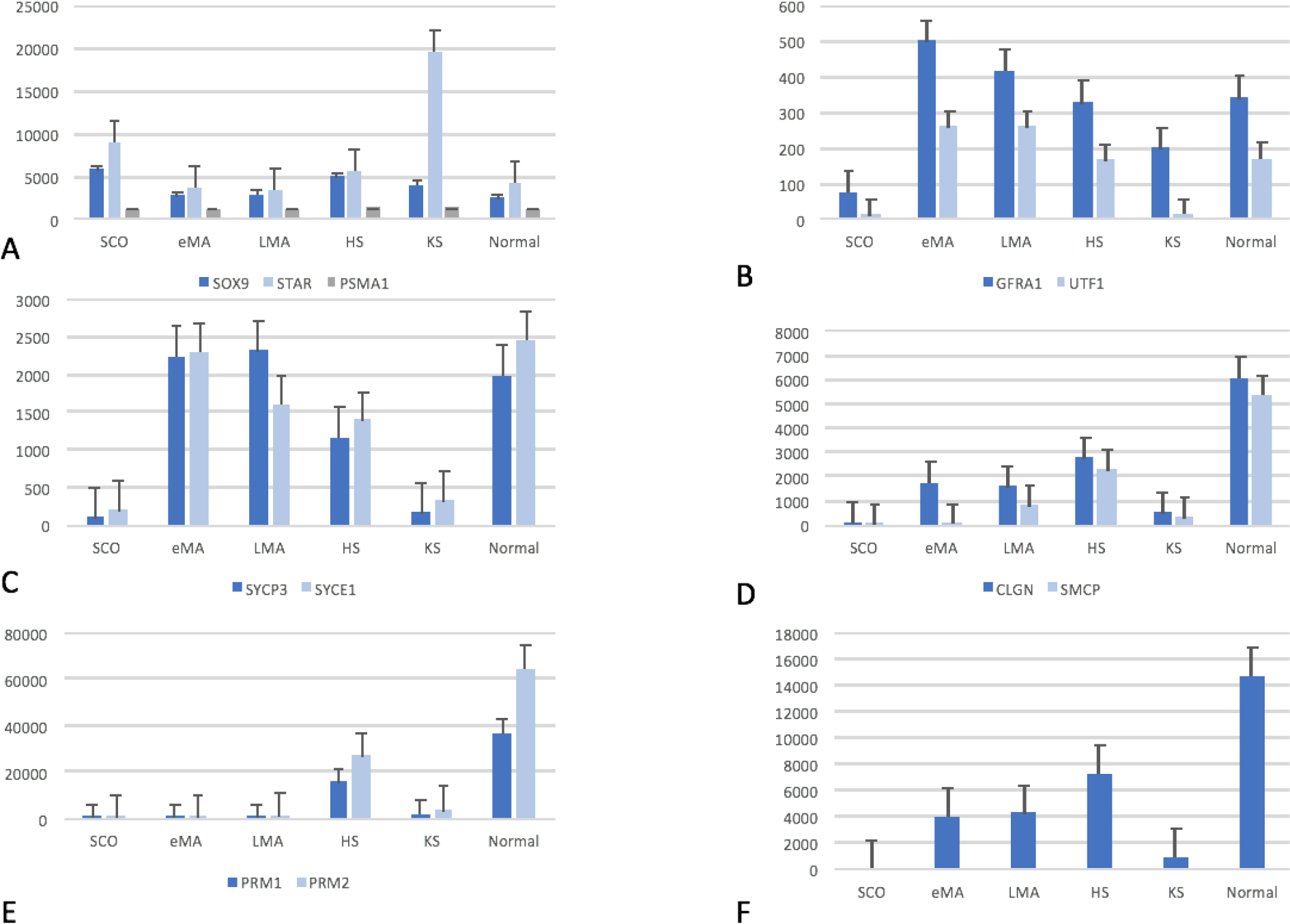
RNASeq normalized transcripts among normal controls and different NOA histologic subclasses: SCO (Sertoli cell only syndrome), eMA (early maturation arrest), NL (normal), HS (hypospermatogenesis), lMA (late maturation arrest), and KS (Klinefelter Syndrome). **A.** Somatic cell transcript markers: SOX9 (Sertoli cells), STAR (Leydig Cells), PSMA1 (Peritubular Myoid Cells), were all comparable between groups. STAR expression was significantly greater among KS testis tissue compared to all other histologic subgroups, in keeping with Leydig cell hyperplasia noted in KS. **B.** Spermatogonial cell markers GFRa1 and UTF1 were significantly down-regulated among SCO and KS in keeping with a general depletion of germ cells as expected. **C.** Meiotic cell markers SYCP3 and SYCE1 were similarly expressed among eMA, lMA, HS and normals, suggesting that similar numbers of meiotic cells containing these transcripts are present. These transcripts were significantly downregulated among KS and SCO. **D.** Post-meiotic cell markers CLGN and SMCP were significantly down regulated in KS and SCO. Only SMCP was significantly down regulated among eMA compared to CLGN suggesting CLGN is transcribed in earlier cells compared to SMCP. Both CLGN and SMCP were significantly down-regulated in LMA compared to normal controls in keeping with the histologic diagnosis of absent haploid germ cells in LMA. **E.** YBX2 bound transcripts PRM1 and PRM2, as identified in animal data, were significantly down-regulated in all conditions compared to normal controls. **F.** YBX2 levels were nearly absent in SCO and KS, but present and down-regulated in eMA, LMA, and HS compared to normal controls.

To verify that the specificity of our detection methods for YBX2 and PRM and to assess the soundness of normalization methods we demonstrated that expression of known markers of Sertoli cells (SCs) SOX9, Leydig cells (LCs) STAR, and peritubular cells PSMA1 was similar in each histological group. There were no statistically significant differences in expression levels of mRNAs for markers for Sertoli cells, Leydig cells, and Peritubular cells, validating the data normalization and ensuring that comparisons were biologically sound.

As the translation of *YBX2* closely follows its transcription in most of the cells and considering that fact that men with MA and normal spermatogenesis have a similar number of meiotic cells, comparable levels of YBX2 protein and PRM1/2 mRNA in normal human testis and men with MA was anticipated. Since SCO testes lack meiotic cells, a lack of *YBX2* expression in men with SCO was expected. However, RNAseq data revealed 2.7x lower expression of *YBX2 mRNA* in men with maturation arrest as compared to controls (p=2.7156E-05). PRM1 and PRM2 are necessary for the exchange of histones to protamines in round spermatids. As expected, *PRM1/2*, were expressed 3,795x and 3,909x less in SCO (p= 2.9676E-07). In keeping with histopathology of late MA, the expression of PRM1/2 was 897 x less in MA (2.9676E-07) as compared to fertile controls. To test if lower than expected expression of PRM1/2 mRNA in MA is secondary to the loss of meiotic cells, expression levels of a known marker of spermatocytes, i.e. SYCP3, was compared between groups using the RNAseq data. There was no difference in SYCP3 expression in eMA, lMA, and normal group indicating that the loss of YBX2 was not due to loss of meiotic cells. No differences were found in the expression of PRM1/PRM2 between eMA and lMA groups. The isolated cell expression analysis shows that PRM1 and PRM2 mRNA are expressed in spermatocytes thus we would expect similar, not decreased levels of PRM1/2 in men with MA. The expression of spermatozoa marker SMCP was 4,129 x lower in SCO (p=4.47E-08) and 173x lower in MA (p=4.08E-08) than in controls as anticipated. Germ cell subtypes were visually confirmed using nuclear and morphologic cellular features from both normal controls and lMA, Figure 2.

**Figure 2.**
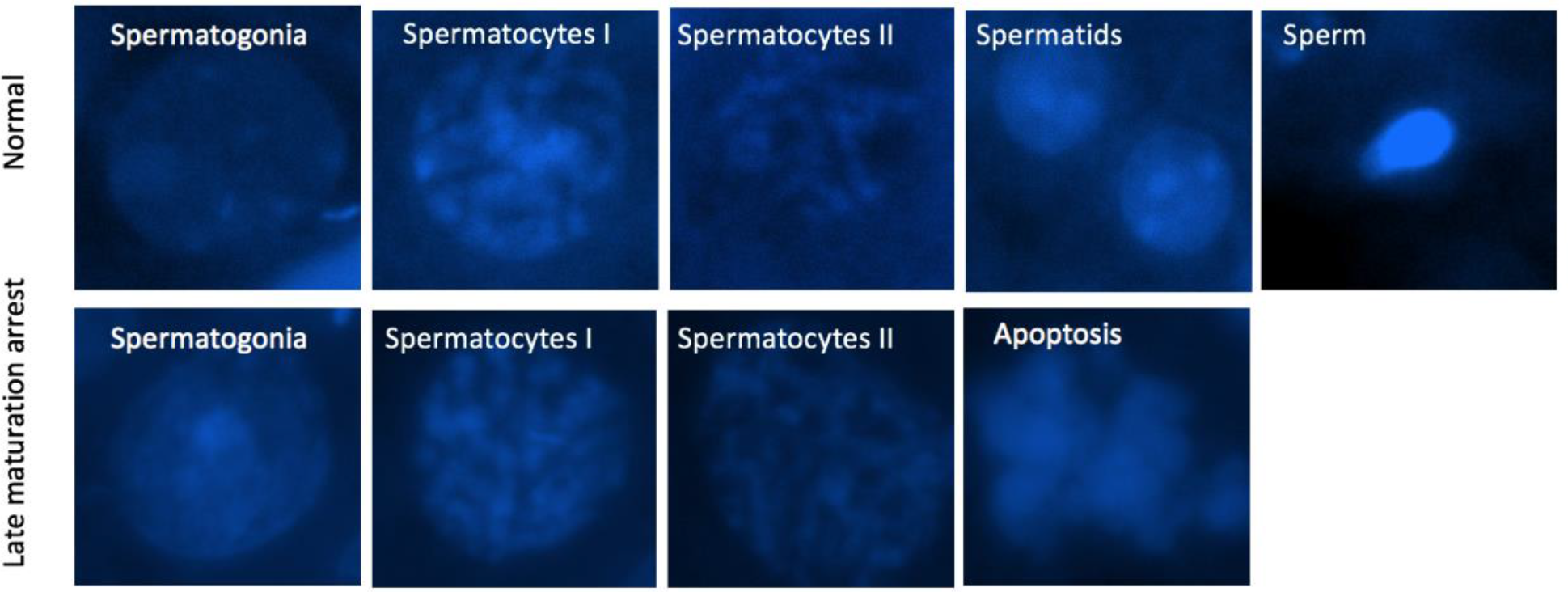
Identifiable cell types in normal and late maturation arrest as determined by DAPI nuclear staining of testis cells from immunofluorescent microscopy slides. Top row (normal spermatogenesis) demonstrates identifiable nuclear features consistent with spermatogonia, spermatocytes, round spermatids and sperm. In the bottom row (late maturation arrest), spermatogonia and spermatocytes are identified as in normal; however, apoptotic spermatocytes are also identified. No round spermatids or sperm were identified in our sections of late maturation arrest. 1000x optical magnification with additional digital enhancement.

As anticipated, lMA was characterized as having meiotic cells present with the loss of round spermatids. Hence our RNAseq showed unexpected loss of YBX2 and PRM1/2 mRNAs in men with maturation arrest who do have meiotic cells in similar numbers of as men with normal spermatogenesis.

To confirm the results of RNAseq and to better understand changes in expression of the YBX2 protein in the human testis, expression of YBX2 using immunofluorescent microscopy in testicular samples from men with normal spermatogenesis and maturation arrest was performed.

YBX2 localized within seminiferous tubules in spermatocytes, Figure 3A. Further analysis of de-convoluted images showed highly granular, cytoplasmic localization of YBX2 in the spermatocytes. High resolution of morphometry of YBX2 positive cells among normal samples reveals at least two subpopulations based upon the size and fluorescent intensity of YBX2, Figure 3A.

**Figure 3A.**
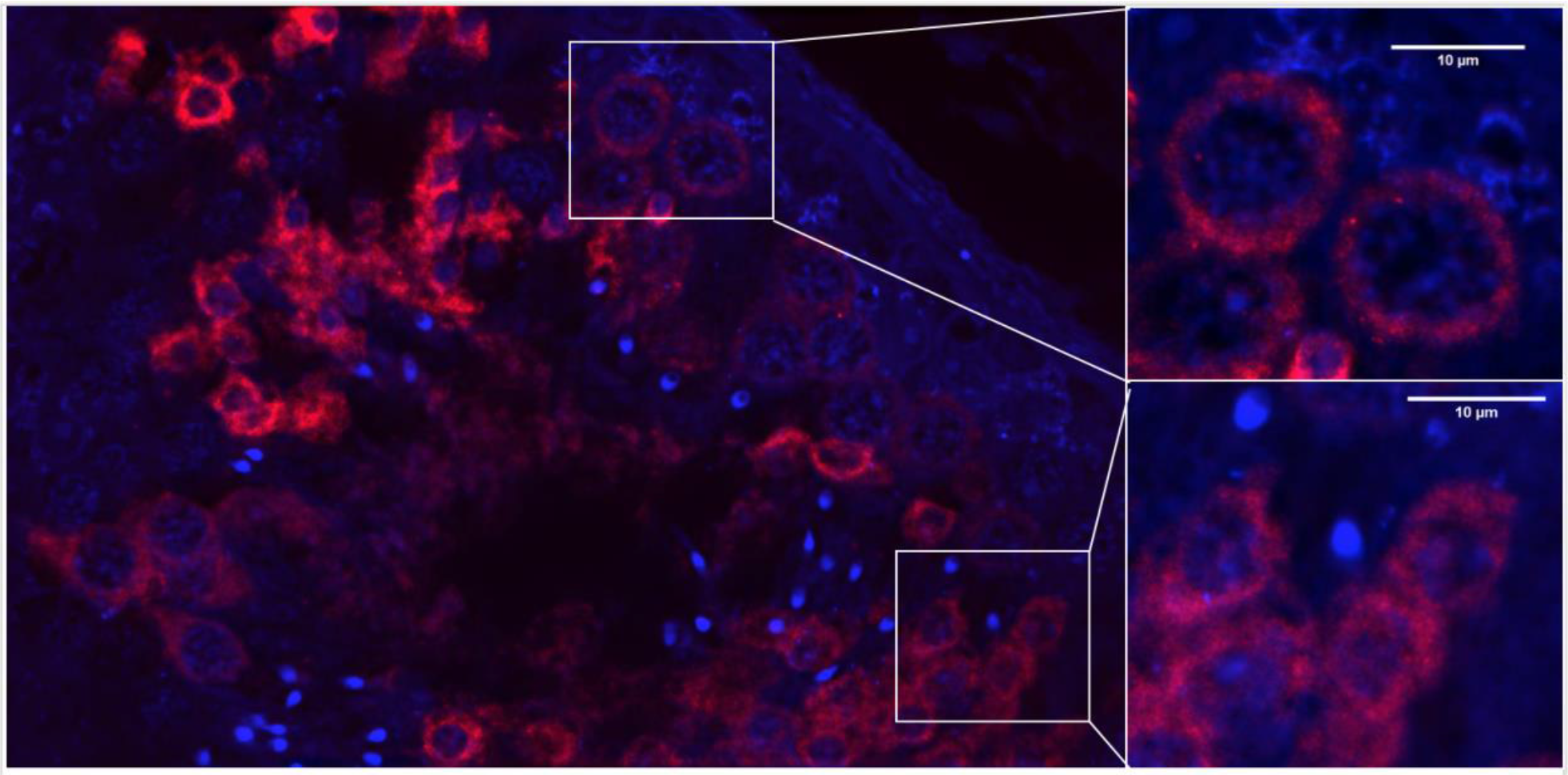
Immunofluorescence analysis of expression of YBX2 protein in normal testis shows bimodal distribution of staining. De-convoluted images taken at 1000x magnification of YBX2 immunofluorescence, with digital zoom of YBX2 positive staining in the cytoplasm of spermatocytes (upper) and round spermatids (lower).

The populations differed concerning both cell size (p <0.001) and nuclear size (p<0.001), the larger cells also demonstrated a larger nucleus but decreased the intensity of YBX2 protein expression levels. Smaller cells showed the higher fluorescent intensity of YBX2 (p<0.001). To further confirm our observation about bimodal staining distribution of YBX2 staining on IF, we used flow cytometry (FC) to measure the size and expression levels of YBX2 per single cell. Flow cytometry results demonstrate two subpopulations of YBX2 positive cells in normal testis, Figure 3B.

**Figure 3B.**
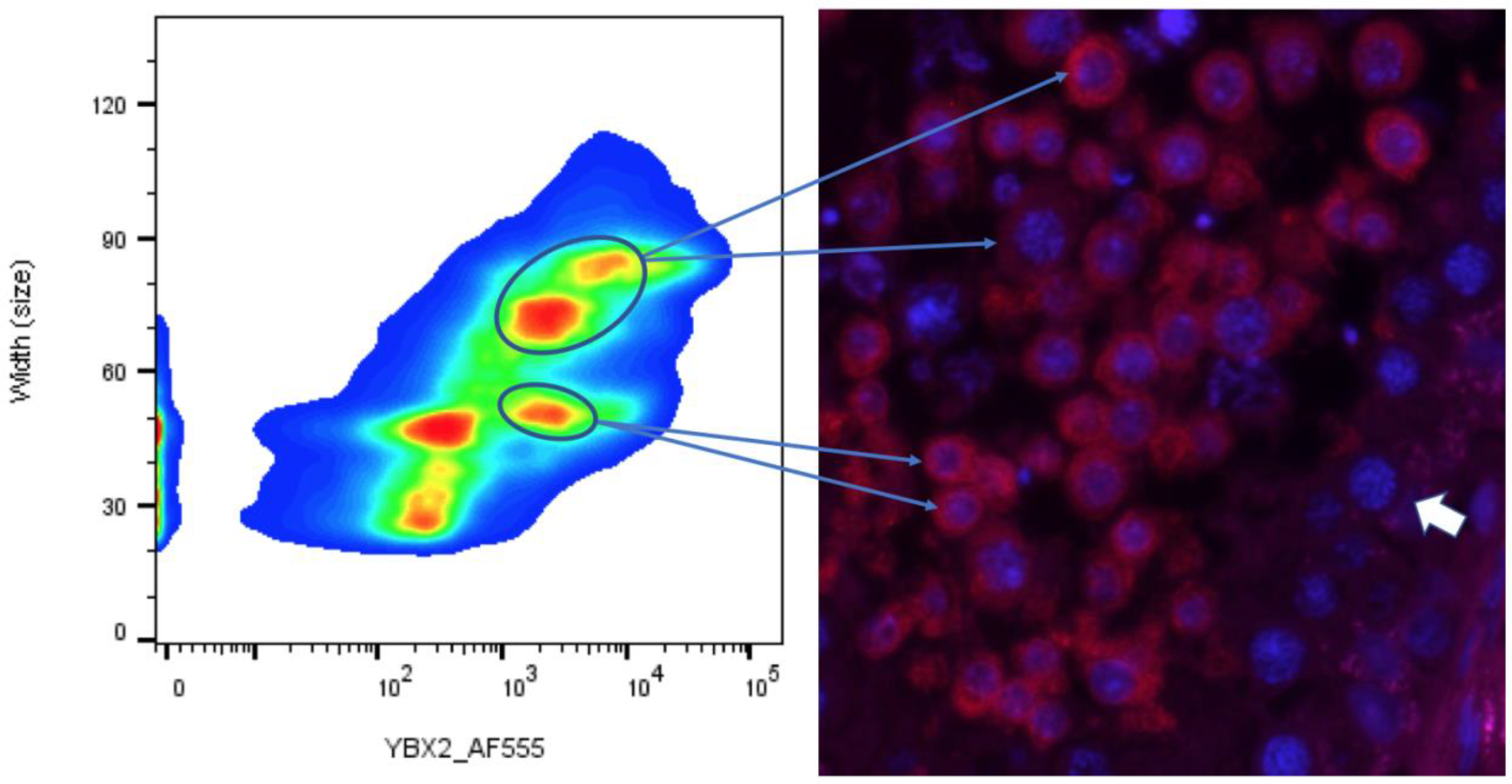
Flow cytometry (left) and immunofluorescence (right) of a normal specimen stained with YBX3 (red on immunofluorescence against blue DAPI nuclear stain). Flow cytometry demonstrates a small population with intense YBX2 staining (bottom right flow cluster), as well as 2 populations of larger YBX2 positively stained cells (upper flow clusters). The white arrow on immunofluorescence demonstrates cells positive for DAPI nuclear stain, but negative YBX2 staining.

Concerning cellular width, the two populations differed in size, 51.7 ± 4.0 vs. 71.4 ± 4.6 (p<0.0001). The smaller cells demonstrated increased fluorescent intensity similar to IF results, 2493 ± 2895 vs. 2138 ± 2374, p<0.0001. More large cells were counted per run compared to small cells (9,483 large cells vs. 6,826 small cells). Our results indicate changes with expression levels of YBX2 during the transition from spermatocyte I to spermatocyte II with a sharp loss of expression of YBX2 when secondary spermatocytes divide into spermatids; this is consistent with the understood mechanism of YBX2 action: retention and sequestration of mRNA which is expressed early but required and translated later in spermatogenesis.

An antibody against SYCP3 protein was used to co-localize SYCP3 and YBX2 within the human testis. Our results indicate that YBX2 is expressed in most but not all SYCP3 positive cells, indicating that SYCP3 may be expressed earlier during spermatogenesis than YBX2, as the spermatocytes progress through spermatogenesis the cells lose SYCP3 and retain YBX2 in latest stages of spermatogenesis, Figure 4A.

**Figure 4A.**
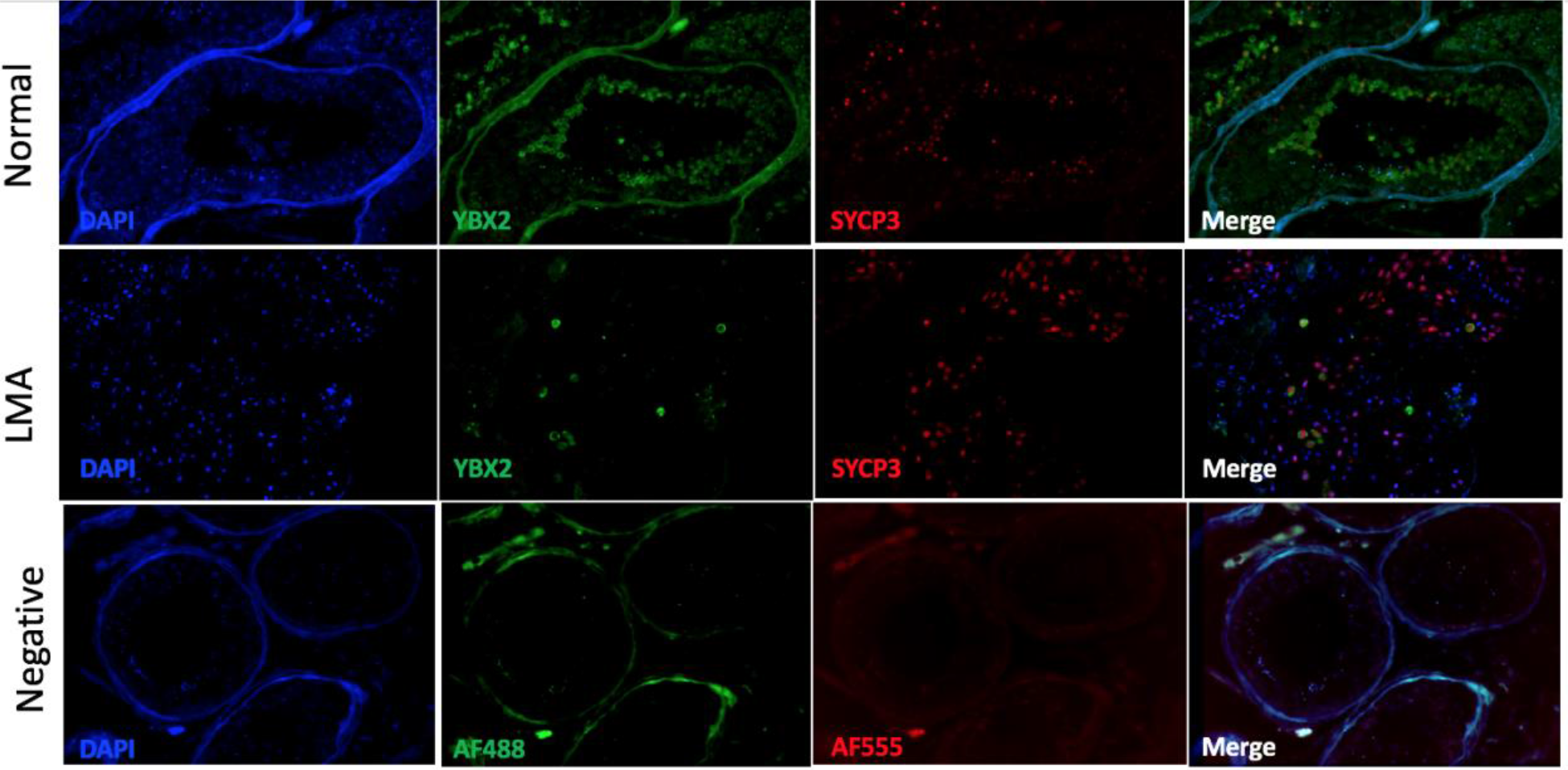
Immunofluorescence images taken at 400x magnification. YBX2 staining localizes to spermatocytes and round spermatids among normal samples. Significant loss of YBX2 is visualized in late maturation arrest (LMA). However, positive staining of SYCP3 among spermatocytes is similar between normal and LMA, suggesting that YBX2 is lost despite the persistence of meiotic cells in LMA. Negative controls do not demonstrate positive staining intra-luminally.

**Figure 4B.**
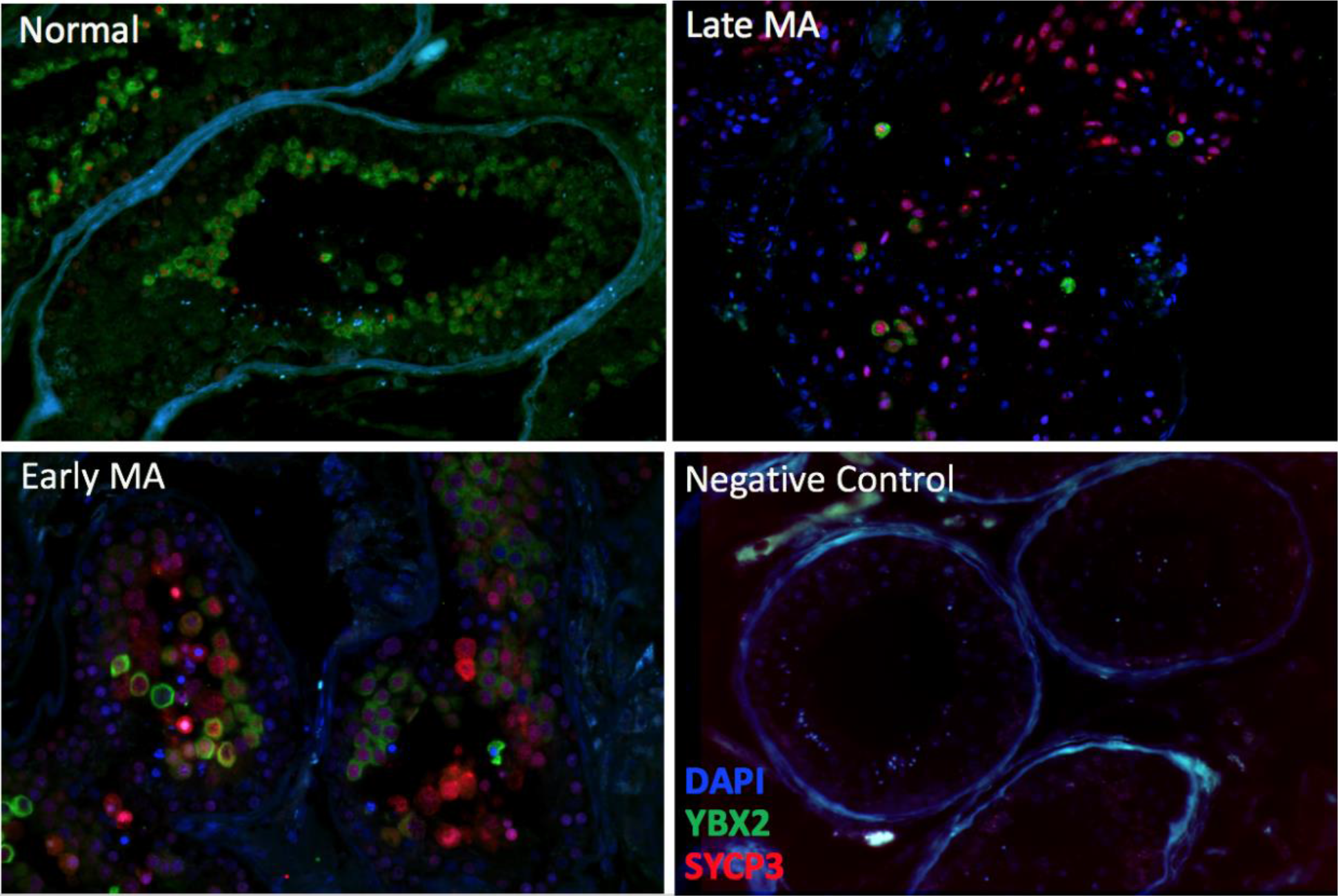
Immunofluorescent 400x magnification of human testis stained for YBX2 (green), SYCP3 (red) and DAPI nuclear stain (blue). A significant number of YBX2 positively stained cells exist in normal spermatogenesis, while very few positively stained cells are present in late MA. However, many YBX2 positively stained cells are observed in early MA suggesting that loss of YBX2 is intricately involved in the pathogenesis of late MA. Negative control contains no primary antibody.

We hypothesize, based upon animal data, that YBX2 is a key ribonuclear protein binding and sequestrating mRNAs needed in later stages of spermatogenesis. Thus, PRM1 and PRM2 expression were assessed in relationship to YBX2, Figure 5A, and 5B. It was clear that in normal testes the loss of YBX2 heralds rapid translation of PRM1 and PRM2 as co-expression of both YBX2 and PRM1 or PRM2 in normal spermatogenesis was not observed in the same cell. Interestingly in LMA PRM2 is sequestrated within the perinuclear area, most likely corresponding to the chromatoid body, Figure 5A and B; thus, indicating that late meiotic arrest may be due to not only the inadequate amount of PRM1/2 mRNA but also abnormal sequestration of PRMs in secondary spermatocytes and lack of trafficking of PRMs from the cytoplasm to nucleus where they function to replace histones.

**Figure 5A.**
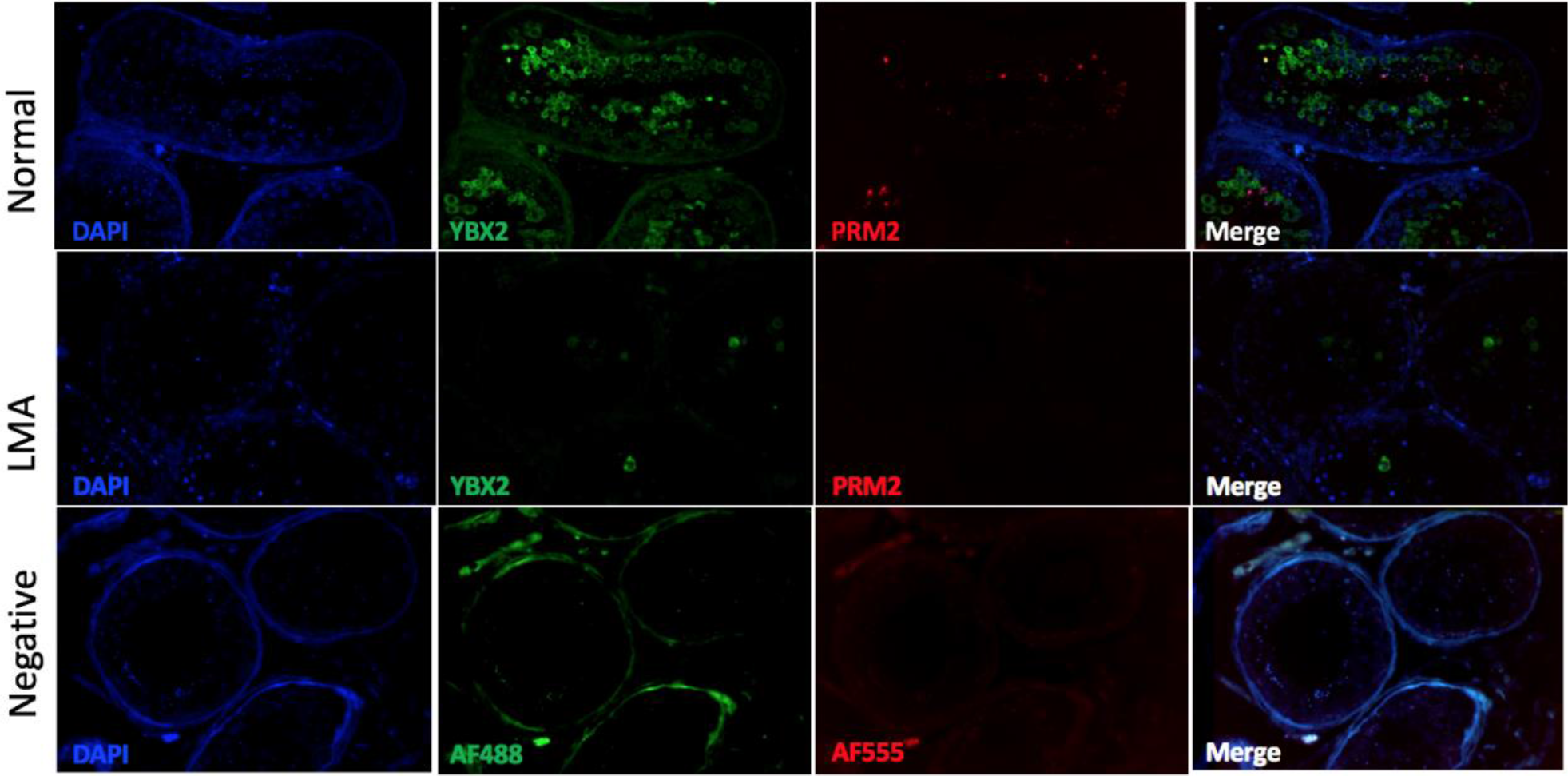
Immunofluorescence images taken at 400x magnification. YBX2 staining localizes to spermatocytes and round spermatids among normal samples. Significant loss of both YBX2 and PRM2 are visualized in LMA. Negative controls do not demonstrate positive staining intra-luminally.

**Figure 5B.**
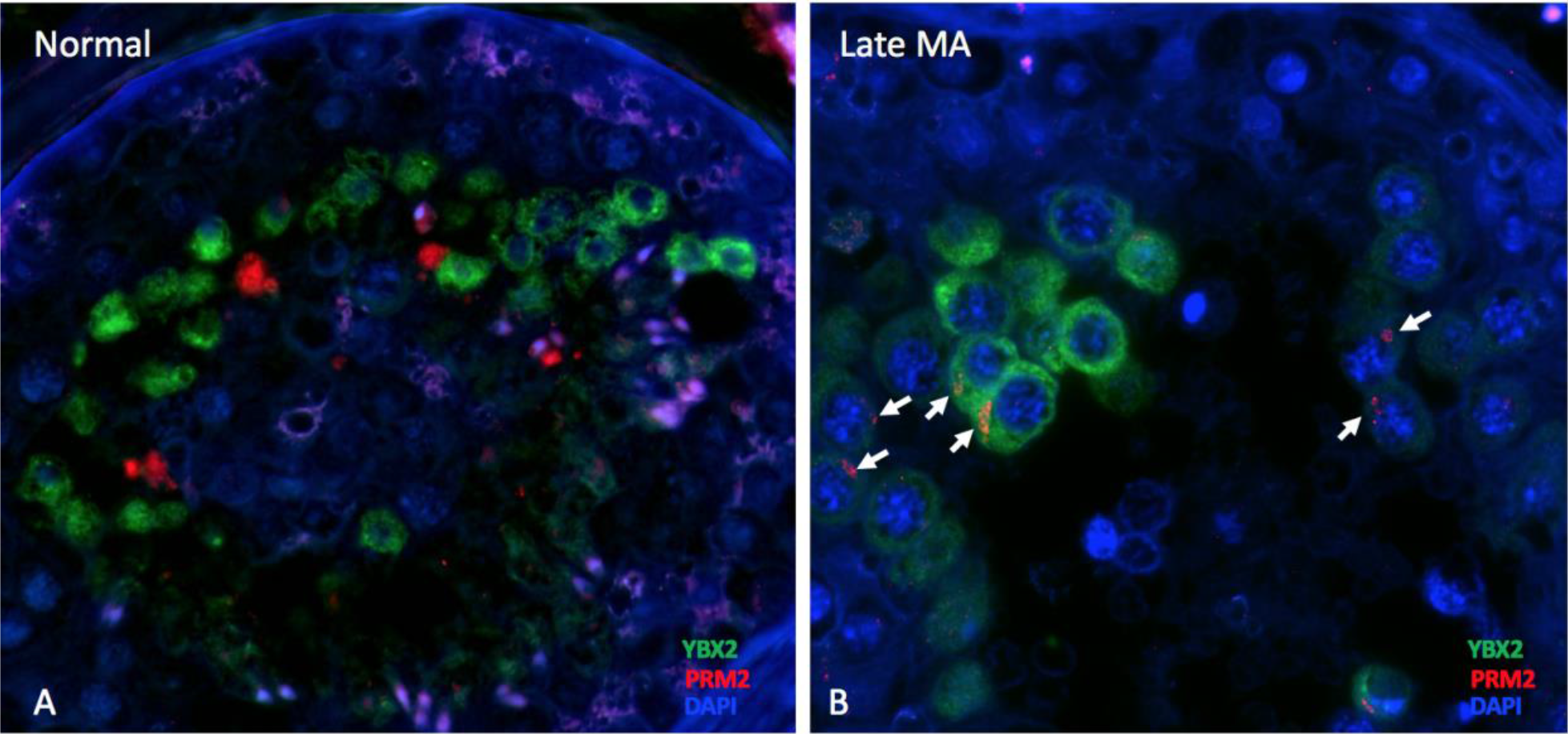
De-convoluted 400x magnification image of YBX2 (green), and PRM2 (red) staining among a human testis biopsy of normal spermatogenesis (A), and late maturation arrest NOA pathology (B). PRM2 staining is limited to spermatids in the biopsy with normal spermatogenesis, separate from YBX2 positively stained cells. However, among late MA, PRM2 protein was identified in the perinuclear region of YBX2 positively stained cells (white arrows), suggesting ineffective translational suppression resulting in premature translation of PRM2 and subsequent inability to complete spermatogenesis.

When YBX2 expression was compared among late and early MA testes, it was clear that despite a normal number of meiotic spermatocytes (SYCP3 positive cells) in lMA, most meiotic cells are not expressing YBX2. This observation confirming that the loss of YBX2 mRNA expression in MA observed on RNAseq is due to actual loss of RNA expression and not the loss of meiotic cell population, Figure 4A and B. YBX2 demonstrated robust staining among spermatocytes and round spermatids in normal human testis. However, among late MA sections with spermatocytes present, YBX2 staining was much less frequent with only 7.9 YBX2 positive cells per tubule compared to 65.1 positive cells per seminiferous tubule among normal tissue (p<0.001). No expression of PRM1 and PRM2 protein was observed in the late maturation arrest other than rare areas of PRM2 sequestration in perinuclear space, Figure 5A and B. This was not expected as PRM1/2 mRNA expression starts early during meiosis thus one would expect PRM1/2 to be present in MA. The sequestration of PRM protein is an intriguing preliminary observation, which will need to be further explored.

Considering that the mRNA levels of YBX2 in eMA and lMA are similar, even adjusted for number of meiotic, Sertoli and Leydig cells, we were surprised to find that the number of YBX2 positive cells in eMA was similar to that of normal controls, but the number of YBX2 positive meiotic cells was drastically reduced in late maturation arrest as compared to normal controls, Figure 4B. To assure that the observed loss of YBX2 expression among meiotic cells in LMA, we verified that number of SYCP3 positive cells were similar in NL, eMA, and LMA. Thus, indicating that loss of YBX2 in LMA is due to abnormal regulation of its expression and not due to sole loss of population.

To better understand the discrepancy in the expression of YBX2 protein among men with normal spermatogenesis, and early and late maturation arrest we measured the expression of transcription factors in all 54 samples and created a multifactorial model to identify the transcription factors which are driving changes in expression of YBX2 between different histological groups. FANTOM5 identified COMP, CDK1, CTCFL, EZH2, KDM5A, PPARG, CTBP1, E2F4, and USF1 as promotor regulators of YBX2. Multifactorial regression analysis demonstrated that COMP levels were significantly lower in eMA and lMA compared to normal controls (p<0.0001). No additional YBX2 promoter regulators: CTCFL, EZH2, KDM5A, PPARG, CTBP1, E2F4, and USF1 were statistically different. Figure 6, demonstrates the expression of each YBX2 promotor among MA and normal controls.

**Figure 6.**
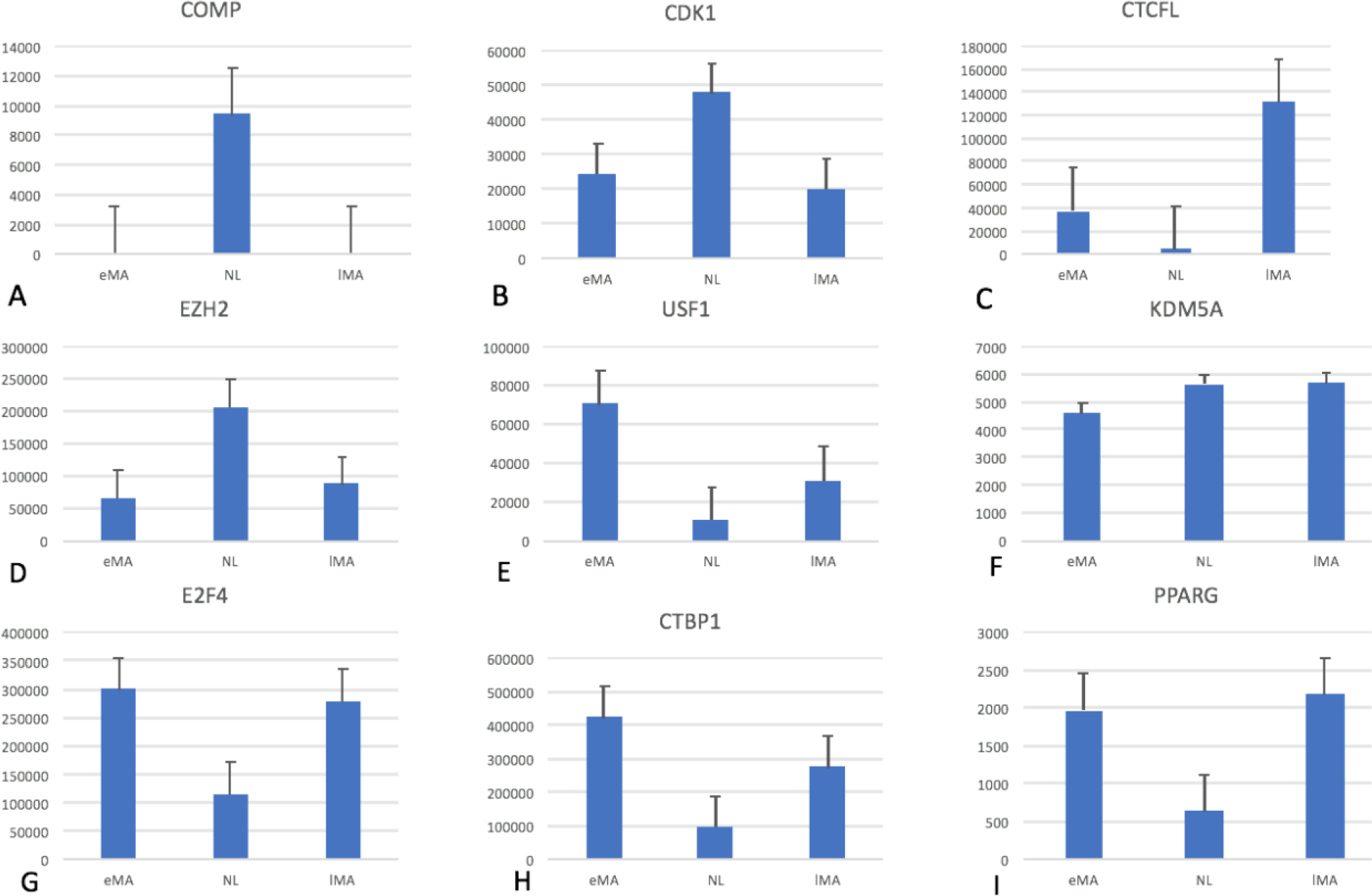
RNA transcript reads for regulators of YBX2 as identified by FANTOM5. Reads are presented among normal controls, as well as early and late maturation arrest. **A)** Expression of COMP is significantly reduced among eMA and lMA patients compared to normal controls. Expression of: **B.** CDK1, **C.** CTCFL, **D.** EZH2, **E.** USF1, **F.** KDM5A, **G.** E2F4, **H.** CTBP1, **I.** PPARG are not significantly different among MA and NL.

**Table 1.** Results of RNA sequencing per histologic subtype of NOA patients: Sertoli cell-only syndrome (SCOS), early maturation arrest (eMA), late maturation arrest (lMA), hypospermatogenesis (HS), Klinefelter Syndrome (KS), and normal controls. Transcripts are shown for spermatocyte markers (SYCP3), and spermatids (CLGN) to compare estimated ratios of cells among histologic subgroups. YBX2, PRM1, and PRM2 are genes of interest and COMP, CDK1, CTCFL, EZH2, KDM5A, PPARG, CTBP1, E2F4, and USF1 were YBX2 regulators identified by Phantom5.

|  |  | Mean Number of Transcripts Per Histologic Subtype |  |  |  |  |  |
| --- | --- | --- | --- | --- | --- | --- | --- |
| Gene Association | Gene | SCO | eMA | IMA | HS | KS | Normal |
| Spermatocyte | SYCP3 | 4.11 | 8215002.08 | 15262305.1 | 177055.85 | 718.75 | 6391713.24 |
| Spermatids | CLGN | 0.63 | 4.68 | 4.66 | 4.89 | 2.93 | 6.96 |
| Translation Repressor | YBX2 | -0.94 | 6.90 | 7.15 | 6.98 | 4.69 | 9.11 |
|  | PRM1 | 1.66 | 2.61 | 3.04 | 10.44 | 7.82 | 12.78 |
|  | PRM2 | 0.51 | 2.27 | 3.56 | 9.97 | 7.48 | 12.39 |
| YBX2 Regulators | COMP | 0.10 | 1.91 | 1.90 | 2.26 | 0.58 | 3.97 |
|  | CDK1 | -0.37 | 4.39 | 4.30 | 3.06 | 1.45 | 4.68 |
|  | CTCFL | -3.79 | 4.58 | 5.12 | 2.38 | -0.43 | 3.63 |
|  | EZH2 | 1.15 | 4.82 | 4.94 | 3.95 | 2.40 | 5.31 |
|  | KDM5A | 3.02 | 3.66 | 3.75 | 3.26 | 2.72 | 3.75 |
|  | PPARG | 0.05 | 3.29 | 3.34 | 2.05 | 0.66 | 2.81 |
|  | CTBP1 | 6.17 | 5.63 | 5.44 | 5.64 | 6.09 | 4.99 |
|  | E2F4 | 5.35 | 5.47 | 5.45 | 5.23 | 5.35 | 5.06 |
|  | USF1 | 5.57 | 4.85 | 4.48 | 4.89 | 5.41 | 4.01 |

In short, YBX2 is expressed in spermatocytes and round spermatids in normal testis. Among men with late maturation arrest, YBX2 expression is significantly down-regulated in meiotic cells at mRNA and protein levels and as such loss of protamines to complete spermatogenesis. In early maturation arrest, the number of YBX2 positive cells appears similar to that found in normal spermatogenesis. Considering that the quantity of YBX2 RNA in late and early maturation arrest is the same, we propose that translation of YBX2 in late maturation arrest is abnormal and suppressed, by yet not clearly understood mechanisms; thus, leading to failure to retain adequate amounts of protamine 1 and 2 mRNA until translation of PRM1/2 can occur in spermatids. Our finding of differences in expression of YBX2 in eMA and lMA indicates that early and late maturation arrest are the result of very different molecular mechanisms.

## Discussion

During spermatogenesis periods of suppression of transcription are needed to complete meiosis. Thus temporal and spacial uncoupling of transcription from the translation is critical to complete spermatogenesis. Small non-coding RNAs can achieve translational control, in addition to nuclear retention of mRNAs, or binding of mRNAs to ribonuclear proteins which attachment of mRNA translation proteins to polyA tail. In the current manuscript, we have focused on understanding the role of YBX2 in maturation arrest and evaluate if loss of YBX2 mRNA expression leads to, or is associated with male infertility.

Using RNAseq, immunofluorescence, and flow cytometry we determined that YBX2 mRNA and protein expression are significantly reduced in patients with late maturation arrest compared to healthy controls in spite of the presence of a similar number of meiotic cells in MA and NL controls.

YBX2 localized predominantly to spermatocytes and round spermatids in normal testes and demonstrated significant loss of detectable protein in round spermatids due to the loss of these cells in MA; however, YBX2 loss also occurred in the corresponding spermatocytes among tissue derived from late maturation arrest, but not to the same degree in early maturation arrest suggesting different molecular underpinnings. Down-regulation of YBX2 in a subpopulation of meiotic cells in late maturation arrest may result in less transcriptional activation, and more critically, the loss of binding capacity for cytoplasmic RNA which are needed in later stages of spermatogenesis; thus, if there is conserved function between murine models and humans, critical genes required for subsequent translation such as PRM1 and 2 will be down-regulated resulting in an meiotic arrest of spermatogenesis as we observed.

YBX2 is necessary for structural integration with RNP complexes, and binding to YRS leading to induction of PRM1 and PRM2 transcription and then PRM mRNA retention, functionally resulting in repression of mRNA translation^17,18^. This process is required among round spermatids in animals, and failure of translational repression results in failed spermatogenesis. This is because post-meiotic regulation requires precise temporal control of transcription and translation as well as appropriate translational suppression. Transcription necessary for post-meiotic differentiation starts early during meiosis and then intensifies among round spermatids^19^. However, the transcripts are translationally repressed for up to 7 days^20,21^. In humans, this may occur earlier as we identified aberrant PRM2 protein adjacent to the nuclear membrane among late meiotic cells in tissue from late maturation arrest. Thus, this may suggest that PRM2 is transcribed and translated during the conclusion of meiosis and successfully sequestered in normal spermatogenesis. The mechanisms of RNA sequestration have not been fully elucidated; however, mRNA is sequestered in the cytoplasm via binding to phosphorylated RNA binding proteins (RBP) such as YBX2 and messenger ribonucleoproteins (mRNP)^22,23^. YBX2 has been determined to have 7 casein kinase two phosphorylation sites and three putative protein kinase C phosphorylation sites suggesting a potential functional regulation via phosphorylation as proposed for other RBPs^12^. mRNP mediated translational suppression appears to be performed globally with little mRNA sequence specificity^24^. Subsequent modification of the mRNPs results in translational activation and is associated with spermatid differentiation^14^. Thus, our observations of down-regulated YBX2 protein expression among meiotic cells in late MA may suggest abnormal YBX2 translational regulation and post-transcriptional processing. These mechanisms have not been studied to date.

Various RNA binding protein (RBP) families have been identified in germ cells^25–27^, including the family in which YBX2 belongs to, Y-Box proteins. This family of proteins includes YBX1, YBX2, and YBX3 (previously termed MSY1, MSY2 and MSY4 respectively)^25–27^. These proteins are evolutionarily conserved across species from insects to higher vertebrates^27^. They bind both RNA and DNA and carry specific functions in different areas of the cell. In the cytoplasm, they bind RNP complexes and function to stabilize mRNA’s and repress translation in maturing germ cells^27,28^. Translational suppression is likely mediated through preferential binding of both YBX2 and to RNA Y-box protein recognition sequence (YRS) ([UAC][CA]CA[UC]C[ACU]) which is implicated in the translational regulation of PRM1, and can repress translation of mRNA^17,18^. YBX2 protein has also been shown to bind small noncoding RNA’s, which often regulate gene expression^14^. On the other hand, two highly abundant miRNAs in testis (miRNA 34c/449c) bind to 3UTR regions of YBX1 and YBX2, thus suppressing the expression of YBX family. In the nucleus, YBX proteins have a role in regulating transcription, splicing, DNA transport and repair^29^. No studies to date have systematically addressed the transcriptional regulation that YBX2 exerts among human testis cells, even as YBX genes play an essential role in regulating genes critical to later stages of spermatogenesis as observed in murine studies. YBX2 has previously been shown to bind RNA in both sequence-dependent and independent manners^19^; thus, it is possible that YBX2 may play both global and mRNA specific translational control at varying time points during spermatogenesis. The impact of YBX2 function is likely significant, since it comprises a staggering 0.7% of all spermatid protein^6^. The YBX2 expression is testis-specific and has been shown to be greatest among round spermatids among mice, where translation is repressed^7,12^. Our findings in human tissue suggest that YBX2 expression initiates earlier in meiotic SYCP3 positive cells and transcends several states of cellular differentiation through to round spermatids as evidenced by our immunofluorescent and flow cytometry data. However, YBX2 protein expression does not localize to spermatogonia in normal or maturation arrest conditions. Questions that remain unanswered include the function of YBX2 changes at different stages of cellular differentiation concerning both function as a transcriptional factor or cytoplasmic RNA binding protein. The state of phosphorylation of YBX2 may alter protein function and potentially result in maturation arrest.

Further evidence supporting the significance of our observations include YBX2 animal knockouts that result in male infertility^10^. Here, relocation of transcripts from within translationally inactive RNA-mRNP complexes to active translational transcripts associated with polysomes^11,19^ was observed. We believe this is demonstrated in our study, which is evidenced by perinuclear PRM2 protein staining in YBX2 positive cells which appears to be prematurely translated and then retained due to loss of translational repression in MA as compared to the normal testis.

Although we convincingly showed that loss of YBX2 expression in meiotic cells is linked to late MA, the regulation of YBX2 is, however, under investigation. In this study, FANTOM5 software was used to predict potential transcription factors that regulate YBX2 expression. COMP which usually induces expression of YBX2 was significantly downregulated among men with MA compared to normal controls. COMP encodes a 524kDa soluble glycoprotein and is expressed in numerous tissues. It has been shown to play a role in collagen organization, and identified as a biomarker in breast cancer, and has been shown to affect genes involved in oxidative phosphorylation, protein processing in the endoplasmic reticulum, and is protective against ER stress-mediated apoptosis among breast cancer cell lines^30^. Thus, lack of COMP may serve as an upstream factor of YBX2 leading to transcriptional down-regulation in MA. The regulation of YBX2 and YBX1 by miRNA, another highly likely mechanism of YBX expression regulation will be reported in a separate paper by our group, as current paper focused on providing evidence if suppression of YBX2 expression is associated with non-obstructive azoospermia.

In summary, we have demonstrated that YBX2 localizes to at least two unique meiotic and post-meiotic germ cell populations in human testis tissue with normal histology. YBX2 mRNA levels are significantly reduced in both early and late maturation arrest compared to normal testis tissue; however, YBX2 protein expression is mostly lost in late maturation arrest compared to both early MA and normal tissue samples despite numerous SYCP3 positive meiotic cells in late MA. Thus, we hypothesize that dysregulation of YBX2 expression with associated loss of availability of protamine mRNA to complete spermatogenesis is an important mechanism of late maturation arrest in humans. We have identified COMP transcription factor as one of the aberrant regulators of YBX2 transcription in MA. Further studies are required to determine the genes under YBX2 control in the nucleus; how post-transcriptional modifications of YBX2 mRNA control its fate, as well as the cell-stage specific RNAs bound to YBX2 in both fertile and infertile men.

## Conclusion

The loss of YBX2 protein expression in men with late maturation arrest is due to loss of YBX2 mRNA and abnormal translational processing yet to be studied. Loss of YBX2 leads to the release of translational suppression with loss of mRNAs sequestrated by YBX2 i.e., PRM1 and 2. The decrease in YBX2 expression is associated with loss of COMP transcription factor. Our findings have significant therapeutic implications; as modulating YBX2 via transcription factors or small non-coding RNAs could potentially restore PRM1 and 2 expression and potentially restore completion of spermatogenesis.

## Acknowledgments

We would like to thank Paula Cohen Ph.D., Andrew Grimson, Ph.D., and Jennifer Grenier, Ph.D., William Wright Ph.D., and John Schimenti, Ph.D. for their constructive feedback during this project.

## Abbreviations

eMA =: early maturation arrest lMA = late maturation arrest
SCO =: Sertoli cell only syndrome
HS =: hypospermatogenesis
PRM1 =: protamine 1
PRM2 =: protamine 2
SMCP =: sperm mitochondria associated cysteine protein
YBX2 =: Y-box binding protein 2
YBX3 =: Y-box binding protein 3
CTCFL =: CCCTC-binding factor like
EZH2 =: enhancer of zeste 2 plycomb repressive complex 2 subunit USF1 = upstream transcription factor 1
KDM5A =: lysine demethylase 5A
E2F4 =: E2F transcription factor 4
CTBP1 =: c-terminal binding protein 1
PPARG =: peroxisome proliferator activated receptor gamma
FANTOM =: functional annotation of the mammalian genome

## Reference

1. Wosnitzer M, Goldstein M, Hardy MP. Review of Azoospermia. Spermatogenesis. 2014;4:e28218.

2. Oliva R, Bazett-Jones D, Mezquita C, Dixon GH. Factors affecting nucleosome disassembly by protamines in vitro. Histone hyperacetylation and chromatin structure, time dependence, and the size of the sperm nuclear proteins.J Biol Chem. 1987;262(35):17016–17025.

3. Steger K. Transcriptional and translational regulation of gene expression in haploid spermatids. Anat Embryol (Berl).1999;199(6):471–487.

4. Hecht NB. Regulation of ‘haploid expressed genes’ in male germ cells. J Reprod Fertil.1990;88(2):679–693.

5. Dadoune JP. The nuclear status of human sperm cells. Micron. 1995;26(4):323–345.

6. Yang J, Medvedev S, Reddi PP, Schultz RM, Hecht NB. The DNA/RNA-binding protein MSY2 marks specific transcripts for cytoplasmic storage in mouse male germ cells. Proc Natl Acad Sci U S A. 2005;102(5):1513–1518.

7. Oko R, Korley R, Murray MT, Hecht NB, Hermo L. Germ cell-specific DNA and RNA binding proteins p48/52 are expressed at specific stages of male germ cell development and are present in the chromatoid body. Mol Reprod Dev.1996;44(1):1–13.

8. Nikolajczyk BS, Murray MT, Hecht NB. A mouse homologue of the Xenopus germ cell-specific ribonucleic acid/deoxyribonucleic acid-binding proteins p54/p56 interacts with the protamine 2 promoter. Biol Reprod.1995;52(3):524–530.

9. Yiu GK, Murray MT, Hecht NB. Deoxyribonucleic acid-protein interactions associated with transcriptional initiation of the mouse testis-specific cytochrome c gene. Biol Reprod.1997;56(6):1439–1449.

10. Yang J, Medvedev S, Yu J, et al. Absence of the DNA-/RNA-binding protein MSY2 results in male and female infertility. Proc Natl Acad Sci U S A. 2005;102(16):5755–5760.

11. Yang J, Morales CR, Medvedev S, Schultz RM, Hecht NB. In the absence of the mouse DNA/RNA-binding protein MSY2, messenger RNA instability leads to spermatogenic arrest. Biol Reprod. 2007;76(1):48–54.

12. Tekur S, Pawlak A, Guellaen G, Hecht NB. Contrin, the human homologue of a germ-cell Y-box-binding protein: cloning, expression, and chromosomal localization. J Androl. 1999;20(1):135–144.

13. Najafipour R, Moghbelinejad S, Samimi Hashjin A, Rajaei F, Rashvand Z. Evaluation of mRNA Contents of YBX2 and JHDM2A Genes on Testicular Tissues of Azoospermic Men with Different Classes of Spermatogenesis. Cell J. 2015;17(1):121–128.

14. Hammoud S, Emery BR, Dunn D, Weiss RB, Carrell DT. Sequence alterations in the YBX2 gene are associated with male factor infertility. Fertil Steril. 2009;91(4):1090–1095.

15. Abdullah L, Bondagji N. Histopathological patterns of testicular biopsy in male infertility: A retrospective study from a tertiary care center in the western part of Saudi Arabia.Urol Ann.2011;3(1):19–23.

16. Lizio M, Harshbarger J, Shimoji H, et al. Gateways to the FANTOM5 promoter level mammalian expression atlas. Genome Biol. 2015;16:22.

17. Chowdhury TA, Kleene KC. Identification of potential regulatory elements in the 5’ and 3’ UTRs of 12 translationally regulated mRNAs in mammalian spermatids by comparative genomics. J Androl. 2012;33(2):244–256.

18. Giorgini F, Davies HG, Braun RE. MSY2 and MSY4 bind a conserved sequence in the 3’ untranslated region of protamine 1 mRNA in vitro and in vivo. Mol Cell Biol. 2001;21(20):7010–7019.

19. Snyder E, Soundararajan R, Sharma M, Dearth A, Smith B, Braun RE. Compound Heterozygosity for Y Box Proteins Causes Sterility Due to Loss of Translational Repression. PLoS Genet.2015;11(12):e1005690.

20. Mali P, Kaipia A, Kangasniemi M, et al. Stage-specific expression of nucleoprotein mRNAs during rat and mouse spermiogenesis. Reprod Fertil Dev. 1989;1(4):369–382.

21. Balhorn R, Weston S, Thomas C, Wyrobek AJ. DNA packaging in mouse spermatids. Synthesis of protamine variants and four transition proteins. Exp Cell Res. 1984;150(2):298–308.

22. Kleene KC. Patterns of translational regulation in the mammalian testis. Mol Reprod Dev. 1996;43(2):268–281.

23. Richter JD, Smith LD. Reversible inhibition of translation by Xenopus oocyte-specific proteins. Nature. 1984;309(5966):378–380.

24. Marello K, LaRovere J, Sommerville J. Binding of Xenopus oocyte masking proteins to mRNA sequences. Nucleic Acids Res.1992;20(21):5593–5600.

25. Cooke HJ, Elliott DJ. RNA-binding proteins and human male infertility. Trends Genet. 1997;13(3):87–89.

26. Wolffe AP, Tafuri S, Ranjan M, Familari M. The Y-box factors: a family of nucleic acid binding proteins conserved from Escherichia coli to man. New Biol.1992;4(4):290–298.

27. Wolffe AP. Structural and functional properties of the evolutionarily ancient Y-box family of nucleic acid binding proteins. Bioessays. 1994;16(4):245–251.

28. Ladomery M, Sommerville J. A role for Y-box proteins in cell proliferation. Bioessays. 1995;17(1):9–11.

29. Matsumoto K, Wolffe AP. Gene regulation by Y-box proteins: coupling control of transcription and translation. Trends Cell Biol. 1998;8(8):318–323.

30. Englund E, Bartoschek M, Reitsma B, et al. Cartilage oligomeric matrix protein contributes to the development and metastasis of breast cancer. Oncogene. 2016;35(43):5585–5596.

